# Changes in patterns of neural activity underlie a time-dependent transformation of memory in rats and humans

**DOI:** 10.1101/303248

**Authors:** Melanie J. Sekeres, Gordon Winocur, Morris Moscovitch, John A.E. Anderson, Sara Pishdadian, J. Martin Wojtowicz, Marie St-Laurent, Mary Pat McAndrews, Cheryl Grady

## Abstract

The dynamic process of memory consolidation involves a reorganization of brain regions that support a memory trace over time, but exactly how the network reorganizes as the memory changes remains unclear. We present novel converging evidence from studies of animals (rats) and humans for the time-dependent reorganization and transformation of different types of memory as measured both by behavior and brain activation. We find that context-specific memories in rats, and naturalistic episodic memories in humans, lose precision over time and activity in the hippocampus decreases. If, however, the retrieved memories retain contextual or perceptual detail, the hippocampus is engaged similarly at recent and remote timepoints. As the interval between the timepoint increases, the medial prefrontal cortex is engaged increasingly during memory retrieval, regardless of the context or the amount of retrieved detail. Moreover, these hippocampal-frontal shifts are accompanied by corresponding changes in a network of cortical structures mediating perceptually-detailed as well as less precise, schematic memories. These findings provide cross-species evidence for the crucial interplay between hippocampus and neocortex that reflects changes in memory representation over time and underlies systems consolidation.

## Introduction

Episodic memories in humans can be construed as having two major constituents: 1) central or schematic elements that are critical to the coherence of the event, and 2) perceptual and contextual details that define the event’s specificity and impart experiential quality. Episodic memories are vulnerable to progressive loss of perceptual and contextual details (Tulving, 1972), but the schematic elements are retained as more general memories for long periods of time (Brainerd and Reyna, 2002; Winocur and Moscovitch, 2011). Notwithstanding their many differences with human memory, memories in rodents share common features and undergo similar changes. For example, contextual fear memories, when initially formed in rodents, are specific to the contexts in which they were acquired. As with human episodic memories, over time, they become more general and can be evoked in different environments that retain the general context with few specific details (Winocur et al., 2007; Wiltgen et al., 2007).

It is widely accepted that the hippocampus plays an essential role in the initial acquisition and storage of memories in both animals and humans. There is considerable controversy, however, regarding its role in the long-term reorganization of memories. The traditional view holds that the hippocampus’s role in memory function is time-limited, after which memories become represented in identical form in neocortex (Kim and Fanselow, 1992; Squire and Alvarez, 1995; Squire et al., 2015). A growing body of evidence, however, suggests that cortically-based memories are qualitatively different than the hippocampus-dependent episodic memories (in humans) and context-specific memories (in rodents) (Dudai, 2012; Kandel et al., 2014).

This evidence has given rise to alternative theoretical models that emphasize an enduring role for the hippocampus in mediating human episodic and rodent context-specific memories (Multiple Trace Theory [MTT], Nadel and Moscovitch, 1997), and a transformation process that allows for the representation of schematic or general memories in neocortex (Trace Transformation Theory [TTT], Winocur and Moscovitch, 2011; Sekeres et al., 2017). The medial prefrontal cortex (mPFC, Frankland et al., 2004; Takashima et al., 2006; Ryan et al., 2015; Bonnici and Maquire, 2017) and other brain regions (posterior cingulate cortex [pCC], precuneus, and angular gyrus, Rugg and Vilberg, 2013) have been implicated in this transformation process.

The present study tests crucial predictions that follow from MTT and TTT; specifically, that reorganized patterns of neural activity, in rats and humans, are related to the quality of a memory as it transforms from one that is perceptually detailed or context-specific to a more schematic or context-general version. The research was also designed to examine several related issues, including the degree of hippocampal activation at remote periods when schematic or generalized memories dominate and, conversely, whether structures, such as mPFC, are implicated in memories that remain context-specific and highly detailed. Notwithstanding differences in tests and delay period, a primary objective of the research was to show that common processes and structures are implicated across species.

In Experiment 1, we tested rats on a contextual fear conditioning task and manipulated context to show that the context-specificity of the learned fear response changes over time. Immediate-early gene expression following retrieval at recent and remote time points measured changes in activity patterns in hippocampus and mPFC that correspond to changes in memory expression. A limitation to current rodent models is the inability to investigate dynamic changes in large-scale neural networks supporting memory across repeated retrieval events. Functional neuroimaging in humans overcomes this limitation. Human episodic memory, like contextual fear memory in rodents, is characterized initially by context-specificity. As in rodents, contextual/perceptual details may be lost over time, while general or schematic features are retained. In Experiment 2, we tested memory for film clips in normal human adults at short and long delays using fMRI to image network activity. We obtained results that complemented those of Experiment 1. This translational approach identifies similarities in hippocampus-dependent memories in rats and humans, and provides evidence for a common memory transformation that can be observed across species.

## Materials and Methods

### Experiment 1: The neural basis of context memory transformation in rodents

#### Subjects

Male Long-Evans rats (Charles River, QC), 3 months old at the start of training, served as subjects for the context fear conditioning experiments. Rats were housed in pairs with unlimited access to food and water, and maintained on a 12 hr reversed light cycle (lights off between 0600 −1800 hr). All behavioral testing occurred during the active phase of the dark cycle. Rats were handled daily for 5 days prior to training. All procedures were approved by Trent University’s Animal Care Committee, and conducted in accordance with the guidelines set by the Canadian Council on Animal Care.

#### Apparatus

Context fear conditioning was conducted in a chamber (70 × 31 × 32 cm^3^) with horizontal striped black and white side and back walls, and a clear Plexiglas front wall. A speaker was affixed to the left side wall. The chamber floor consisted of metal bars spaced 1.3 cm apart, and the Plexiglas roof was ventilated to allow air circulation. The conditioning chamber was placed on a table, 1.3 m above the floor, located in a sound-attenuated laboratory testing room (2.9 × 1.9 × 2.7 m), dimly lit with overhead lighting. The conditioning chamber served as Context-A (CXT-A) for memory testing. The novel chamber used for Context-B (CXT-B) (29 × 29 × 31 cm^3^) had four clear Plexiglas walls, a metal grid floor, and a ventilated Plexiglas roof. The chamber was placed on a desk located in a different sound attenuated test room (4.3 × 3.3 × 2.7 m) with standard laboratory furniture (desk, chairs, sink), and lit by a desk lamp. A video camera was mounted on a tripod in front of each test chamber to record freezing behavior during each condition and test session.

#### Behavioral methods

Eight rats were randomly assigned to each of the five conditions: Home cage control, (HC); Short-Delay, CXT-A (SD-A); Short-Delay, CXT-B (SD-B); Long-Delay, CXT-A (LD-A); Long-Delay, CXT-B (LD-B).

#### Context pre-exposure and contextual fear conditioning

Prior to fear conditioning, each rat was individually placed in the conditioning chamber and allowed 30 min to explore. Twenty-four hours later, the rat was transferred back to the test room, placed in the conditioning chamber, and allowed 3 min to explore. Ten tone-shock pairings (tone: 2000 Hz; 90 db, 30 s; shock: 1.5 mA, 1 s) (TechServe, Model 452A shock generator) were then administered with a variable interval (30 s – 120 s) between pairings. After the last shock, freezing behavior was recorded every 8 s for 64 s (8 observations). The rat was then returned to its home cage. The total time spent freezing during each session was also recorded. Freezing behavior was calculated using an 8-second time sampling procedure for each minute during which freezing was assessed (8 observations per minute). Freezing behavior was defined as the absence of any visible movement aside from respiration. The percentage of time spent freezing was calculated by dividing the total number of observed freezing responses by 8 for each minute. Fear conditioning procedures were adapted from Anagnostaras et al. (1999), and have been routinely used in our lab (Winocur et al., 2007; Winocur et al., 2009; Winocur et al., 2013).

#### Context fear testing

Original Context (CXT-A): Either 24 hr (Short-Delay, SD-A) or 30 days (Long-Delay, LD-A) after conditioning, the rat was returned to the test room, and placed in the conditioning chamber for 8 min. No shock was administered during testing. Freezing behavior was recorded every 8 s (64 observations) by the experimenter. The total time spent freezing during the test session was also recorded. The rat was then returned to its home cage.

Novel-Context (CXT-B): Either 24 hr (Short-Delay, SD-B) or 30 days (Long-Delay, LD-B) after conditioning, the rat was brought to the test room and placed in the novel chamber for 8 min. Testing was conducted identically to what is reported for CXT-A. See Figure 1A for a schematic of the study design.

Home Cage (HC) control: The HC group was included to control for baseline expression of the immediate-early gene (IEG) c-Fos. These rats were maintained in their home cages throughout the experiment, and did not undergo any behavioral training or testing. After 30 days, rats were removed from their home cage, sacrificed, perfused, and brains were prepared for immunohistochemistry.

#### Perfusion and histology

Immediately following testing, rats were transferred to a quiet, dark holding room. 90 min following test (or HC) conditions, received an overdose IP injection of sodium pentobarbital, and were intracardially perfused using phosphate-buffered saline (PBS) followed by 4% paraformaldehyde (PFA). Brains were removed from the skull, fixed in PFA for 24 hr at 4°C, then transferred to a PBS and 0.02% sodium azide solution and stored at 4°C until sectioning. Using a vibratome, brains were sectioned coronally (30 μm slices) across the entire anterior-posterior extent of the brain. Five serial sections per well were stored in a PBS and 0.02% sodium azide solution. For each brain, one section per well was randomly sampled across the range of the anterior cingulate cortex (aCC) and hippocampus. For the aCC, 4 – 10 sections ranging between 1.70 mm to – 0.92 mm A/P relative to bregma according to the Paxinos and Watson atlas (1997) were sampled for analysis of c-Fos protein expression. For the hippocampus, 10 – 16 sections ranging between −2.30 mm to −6.04 mm A/P were sampled. 0-3 rats per condition were excluded from c-Fos analysis due to poor tissue integrity resulting in weak immunohistochemical labeling.

#### Immunohistochemistry

Coronal brain sections were washed in PBS, then incubated with rabbit anti-c-Fos polyclonal primary antibody (1:1000, PBS and 0.3% Triton X-100, Calbiochem) at 4°C for 48 hr. Sections were washed 4 × 10 min in PBS, then incubated with donkey-anti-rabbit Alexa 568 secondary antibody (1:200, Molecular Probes) for 2 hr at room temperature. Sections were washed with PBS and then mounted with PermaFluor mounting medium (Thermo Scientific, Fremont, CA) on glass slides, and coverslipped.

#### c-Fos quantification

Stained sections were analyzed using a fluorescent microscope (Nikon, MBA 92010 Eclipse NI). Images were taken at 10X magnification using a digital camera (DS-QiMc-U3), and digitally stitched together using NIS-Elements software (Nikon, version 4.1.3) software to reconstruct each region of interest (hippocampus, aCC). Within each section, hippocampal subregions (CA1, CA3, DG) were outlined for the dorsal and ventral regions, then combined to determine dorsal and ventral hippocampal area. cFos positive nuclei in each subregion were manually counted using ImageJ (RRID:SCR_003070). For the prefrontal cortical sections, the aCC was outlined, and c-Fos positive nuclei were similarly counted. c-Fos is a commonly used marker of neuronal activity (Greenberg and Ziff, 1983). For each sampled section, c-Fos expression was counted bilaterally for the aCC, and unilaterally for the hippocampus. The total number of c-Fos positive cells per region of interest was divided by the outlined area to generate a normalized cells/area value. c-Fos expression was analyzed for home cage (HC) control brains to determine baseline c-Fos expression in each region of interest. Values of c-Fos positive cells/area from each of the four experimental conditions (SD-A, SD-B, LD-A, LD-B) were divided by the HC control values to determine the percent of change in IEG expression for each condition.

Experimental Design and Statistical Analysis A schematic of the experimental timeline can be seen in Figure 1A. ANOVAs were conducted for freezing behavior and on for c-Fos expression measures. All behavioral and histological statistical analyses were conducted using SPSS 23 (RRIS:SCR_002865).

### Experiment 2: The neural basis of episodic memory transformation in humans

#### Participants

Twenty healthy, right-handed participants (12 female), ranging in age from 21-31 years old (mean age 24.05, SD 2.78), were recruited through the participant database at Baycrest. Participants were fluent in English, and screened using a detailed health questionnaire to exclude psychiatric and neurological disorders, previous head injuries, or other health problems and/or medications that might affect cognitive function and brain activity, including strokes and cardiovascular disease. All procedures were approved by Baycrest’s Research Ethics Board, and conducted in accordance with the guidelines set by the Tri-Council Policy Statement: Ethical Conduct for Research Involving Humans. All participants gave written informed consent, and were reimbursed $100 for their participation in the study.

#### Behavioral methods Film clip stimuli

Forty film clips were used to test episodic memory. Film clips have been previously used as naturalistic stimuli for an ecologically-valid memory paradigm (Ben-Yakov and Dudai, 2011; Bird et al., 2015; Furman et al., 2007, 2012; Sekeres et al., 2016; St-Laurent et al., 2014; St-Laurent et al., 2016). Clips were 23 s in duration and were taken from foreign films (i.e. non-English language films) with limited dialogue (the same clips were used in previous studies; St-Laurent et al., 2014, 2016; Sekeres et al., 2016). Two series of 20 clips were equated on four feature categories: visual complexity, story complexity, sound complexity, and emotional content to ensure comparable content of clips used at each test delay (Sekeres et al., 2016). For each participant, the two series of clips were pseudo-randomly assigned for testing either immediately (0d) or 7d after the encoding session.

#### Task

Procedures were based on those developed for a previous study (St-Laurent et al., 2014, 2016). Prior to scanning, participants were read a set of instructions, and then performed a practice session in which they watched two sample clips and performed the memory retrieval task. Participants were told they would be tested on their memory for the clips following varying delays, and instructed not to rehearse the information in the interim. Once in the scanner, they were again briefed on the instructions for the task. All experimental stimuli were viewed through a mirror affixed to the head coil, and responses to the memory ratings were recorded using a button box taped to the right hand. Experimental stimuli were presented using E-Prime 2 (version 2.0.10.242, E-Studio, Psychology Software Tools Inc.).

#### Encoding session in scanner

During encoding, participants viewed the 40 film clips, presented in randomized order. Each clip was given a unique title (i.e. “Boy, Girl and Balloon”) that served as a retrieval cue in the retrieval portion of the experiment. The title appeared centrally on the screen for 4 s immediately before and after the clip played. Clips were centrally presented on a computer screen. Sound was delivered through a rimless Avotech headset. Participants were instructed to pay attention to the title and content of each clip. A fixation cross was presented for 4 s between each clip. Encoding occurred across four runs in the scanner, with 10 clips presented in each run. No response was required during the encoding session. Immediately after the four encoding runs, a 5 min resting state scan was conducted. See Figure 2A and 2B for study timeline and design schematics.

#### Retrieval session in scanner

Memory for a series of 20 clips (see Film Clip Stimuli) was tested either immediately (0d) after the encoding session, or after a 7d delay. Clips were assigned pseudorandomly to a retrieval session timepoint in a manner that was counterbalanced across participants. Participants were presented with the title of a clip for 16 s, during which they were instructed to visualize the clip in their mind, from beginning to end. Next, they used a key pad to rate their memory retrieval for the clip’s story content, on a scale of 1 (no story content) to 4 (all story content). Story content refers to the central plot line of the story (“what happened”), and events central to the progression of the episode (Berntsen, 2002; St-Laurent et al., 2014, 2016; Sekeres et al., 2016). Next, participants rated the vividness of perceptual details retrieved in a similar way (rating of 1 = no perceptual details, rating of 4 = most vivid memory). Perceptual details referred to visual (colors, lighting, textures, facial features, clothing, positions of objects, background details, weather, lighting conditions, etc.) and auditory details (talking, laughing, background music, street sounds). A fixation cross presented centrally on the screen for 4 s separated the retrieval period for each clip.

#### Retrieval session outside scanner

Participants next performed a post-scan test session. During this session, participants were again cued with the title of the clip they had retrieved in the scanner, and asked to verbally report the story content details they recalled while in the scanner (what happened, who did what, what was the situation). Participants were next asked to verbally report, within a maximum of 60s, any perceptual (visual or auditory) details they experienced in their mind’s eye while they recalled the clip in the scanner. Recordings of verbal responses were transcribed and scored according to a system described below. The presentation order of clips was randomized within each retrieval session.

The post-scanning retrieval testing was conducted on a desktop computer using E-Prime 2 in a sound-attenuated room. Recording failed during the verbal retrieval session for one participant, so verbal retrieval data are presented for 19 participants.

#### Scoring and analysis of behavioral data

Self-report ratings of story content and vividness of perceptual details were averaged across clips for each delay. As described above, two separate recordings of the verbal retrieval responses were obtained for each clip to encourage participants to report what they recalled about a clip’s storyline and perceptual content. The recordings were manually transcribed and responses were coded and scored to categorize central elements (indicative of story content) and peripheral details (reflecting perceptual details). Central elements were story details that could not be modified or omitted without changing the plotline of the story (Berntsen, 2002). In order to score central elements consistently, 5 to 6 central story points were identified for each clip and recorded as a ‘central narrative’ (see Sekeres et al., 2016 for a list of central story points for each clip, and for an example of a coded transcript). A participant was given a score of one for each item of retrieved information that corresponded to a point in the central narrative for that clip. Peripheral details were considered any additional descriptive information, including perceptual, and emotional details. One peripheral point was scored for each peripheral story detail reported during the verbal retrieval session. Notably, there was an upper limit to the number of central points a participant could score, but no such limit for peripheral points. To control for the different baseline conditions for each type of detail (central or peripheral), a t-test (2-tailed) of the percentage of details retained (ie. Percent Retained = [(7d/0d) × 100]) between the immediate retrieval (0d) test and the 7d retrieval test was also conducted.

For each clip, both central elements and peripheral details were coded and tallied across the first recording (participant probed for story content) and second recording (participant probed for perceptual details) by an experimenter (S.P.) blind to the delay condition. A subset of recordings was scored by a second experimenter (M.J.S.) to confirm an acceptable rate of 90% inter-rater reliability in detail scoring. Each reported detail was classified as either central or peripheral. No additional points were assigned for repeated details, or for unrelated information about the film clips (i.e. opinions or speculations). Errors in central elements and peripheral details were also calculated. Errors were considered to be any recalled details that did not match the information presented in the film clip. For each type of detail (central or peripheral), the total number of errors was subtracted from the total number of correct details for each clip (i.e. Retrieval Success = # correct details – # errors) to determine the corrected memory retrieval success scores used in the final data analyses. For each participant, the corrected central and peripheral details were averaged across all clips for each delay condition (0d and 7d).

For all behavioral analyses, we excluded ‘forgotten’ retrieval trials, which were trials given memory retrieval ratings of ‘1’ (indicating low memory for story content and low vividness of perceptual details), and for which there were no central elements or peripheral details reported during the verbal retrieval session. We also excluded trials in which the participant reported details corresponding to the wrong film clip. Although these data were excluded in the main analyses, analyses that included these data produced the same pattern of results (data not shown, but available upon request).

Experimental Design and Statistical Analysis

A schematic of the experimental timeline and design can be seen in Figures 2A, 2B. Repeated measures ANOVAs were conducted for the ratings and detailed retrieval measures, and t-tests were conducted to assess differences in the percentage of rating and details retained, and for differences in the number of errors and the number of forgotten trials between 0d and 7d retrieval. Data analysis was performed using SPSS 23. Statistical analyses of brain imaging data are described below.

#### fMRI methods

##### Image acquisition and preprocessing

Participants were scanned using a Siemens Trio 3T scanner. Anatomical scans were acquired with a three-dimensional magnetization-prepared rapid acquisition with gradient echo (MP-RAGE) sequence (repetition time (TR) = 2000 ms, echo time (TE) = 2.6 ms, field of view (FOV) = 256 mm, slice thickness = 1 mm, 160 slices. Functional runs were acquired with an echo planar imaging (EPI) sequence, with 139 volumes for each retrieval run (TR = 2.2s, TE = 27 ms, flip angle = 62°, FOV = 225 mm, 64 × 64 matrix, 36 3.5 mm (skip 0.5 mm) thick axial slices, positioned to image the whole brain. Slices were obtained from an axial oblique orientation, parallel to the Sylvian fissure.

Preprocessing of the image data was performed with Analysis of Functional Neuroimages (AFNI, Cox, 1996). This included regressing out physiological artifact using RETROICOR, rigid motion correction, spatial normalization to Montreal Neurological Institute (MNI) space, and smoothing with an 8 mm Gaussian filter (the final voxel size was 4 × 4 × 4 mm). We also regressed out white matter, cerebral spinal fluid, and vasculature (Campbell et al., 2013; Anderson et al., 2014). As motion has been demonstrated to affect brain-activity measures, even after standard correction procedures (Power et al., 2012), we followed a motion-scrubbing procedure described in Campbell et al., 2013. Briefly, this procedure uses a multivariate technique to identify outliers in both the motion-parameter estimates and fMRI signal itself. Where such outliers co-occurred (never more than 5% of the total volumes), we removed the fMRI volumes and replaced them with values interpolated with cubic splines. This method has the advantage of suppressing spikes, yet keeping the length of the time course intact across subjects.

#### Partial-Least-Squares analysis

The image data were analyzed with Partial Least Squares (PLS; McIntosh et al., 1996; McIntosh and Lobaugh, 2004), a multivariate analysis technique that identifies whole-brain patterns of covariance related to the experimental design (task PLS) in a single step for multiple groups. This method is similar to principal component analysis (PCA), in that it identifies a set of principal components, or “latent variables” (LVs), which optimally capture the covariance between two sets of measurements (Friston et al., 1993). PLS uses singular value decomposition in a data-driven approach to reduce the complexity of the dataset into orthogonal LVs that attempt to explain the maximum amount of covariance between the task conditions and the BOLD signal. In task PLS, each brain voxel has a weight, known as a salience, indicating how strongly that voxel contributes to the LV overall. The significance of each LV as a whole was determined with a permutation test (McIntosh et al., 1996) using 1000 permutations. In addition, the reliability of each voxel’s contribution to a particular LV was tested by submitting all saliences to a bootstrap estimation of the standard errors (SEs; Efron, 1981), using 1000 bootstraps. Peak voxels with a salience/SE ratio ≥ 3.0 (p <.001) are considered to be reliable (Sampson et al., 1989).

Clusters containing at least 5 reliable contiguous voxels were extracted, with a local maximum defined as the voxel with a salience/SE ratio higher than any other voxel in a 2 cm cube centered on that voxel (the minimum distance between peaks was 5 mm). Coordinates of these locations are reported in MNI standard coordinate space (Mazziotta et al., 2001). Because the extraction of the LVs and the corresponding brain images is done in a single step, no correction for multiple comparisons is required. Finally, to obtain summary measures of each participant’s expression of each LV spatial pattern, we calculated brain scores by multiplying each voxel’s salience by the BOLD signal in the voxel, and summing over all brain voxels for each participant in each condition. These brain scores were then mean-centered (using the grand mean across all subjects and conditions) and confidence intervals (CIs; 95%) for the mean brain scores in each condition were calculated from the bootstrap. Following procedures used elsewhere (e.g. McIntosh et al., 2004; Garrett et al., 2010; Grady et al., 2010; Anderson et al., 2014), conservative estimates of differences in activity between conditions and between groups were determined by a lack of overlap in these bootstrapped CIs. That is, non-overlapping intervals between conditions within a group, or between groups within a condition, indicated a significant difference.

#### fMRI task analysis

To assess modulations of BOLD activity across the conditions, we first conducted a mean-centered task PLS analysis (blocked design, where each clip was defined as a block) that contrasted the mean activity (averaged over all blocks across runs) in the retrieval task and fixation at the immediate (0d) and 7d delay conditions (Figures 3A, 3C, Table 1). Brain scores associated with each LV are shown in Figures 3B, 3D. We also ran a PLS analysis directly contrasting the 0d and 7d retrieval tasks (Figure 3E, 3F, Table 2). All retrieval analyses contained trials in which participants reported successfully retrieving the story and perceptual content from the clip (ratings of 2, 3, or 4). Forgotten trials (assigned in-scanner ratings of 1s) were excluded from the retrieval analyses. Finally, we contrasted clips that were recalled with high vividness (in-scanner ratings of 3 and 4 for both story content and perceptual details) (Geib et al., 2015) between the 0d and 7d retrieval tasks to determine differences in retrieval activity at each delay when the memory for the clips remained vivid (Figure 4A, 4B, Table 3). The number of vividly retrieved clips at each delay ranged from (7-19 clips for 0d delay, and 1-14 clips for 7d delay). We also ran analyses for the highly vivid trials including only participants with at least 5 (n = 16), at least 6 (n = 13), and at least 7 (n = 10) of the highly vivid trials, and obtained similar results as the analysis that included all subjects (data not shown, but available upon request).

## Results

### Experiment 1: The neural basis of context memory transformation in rodents

#### Behavioral Results: Retrieval of contextual fear memory

We first set out to replicate our previous behavioral findings to demonstrate the reliability of contextual fear generalization over time (Winocur et al., 2007; Wiltgen and Silva, 2007; Einarsson et al., 2014). Freezing behavior, the measure of fear memory, was assessed in the original conditioning context (CXT-A), or in a novel context (CXT-B) following a short delay (SD; 1 day) or a long delay (LD; 30 days). The time course of freezing was assessed over an 8 min test period for each of the four groups. As all groups showed a decline in freezing in the last 2 min of the test, only the first 6 min of the test were analyzed (8 min test data not shown, but available upon request). A 2 × 2 ANOVA assessed freezing behavior, with Context (CXT-A, CXT-B) and Delay (SD, LD) as between-subject factors. Significant main effects were found for Context, with more freezing in CXT-A than in CXT-B (F(_1,31_) = 7.178, p = 0.012, η^2^_p_ = 0.204), and for Delay, with more freezing at the LD than SD (F(_1,31_) = 4.409, p = 0.045, η^2^_p_ = 0.136). A significant Context x Delay interaction (F(_1,31_) = 11.184, p = 0.002, η^2^_p_ = 0.285) confirmed that rats exhibited context-specific memory at the short delay, freezing at higher levels in CXT-A than CXT-B. At the long delay, groups displayed equivalent levels of freezing in both contexts (Figure 1B), consistent with the idea that the memory generalized over time. These effects provide validation for the next stage of the research which aimed to identify underlying patterns of neural activity supporting changes in the quality of context memory over time.

#### c-Fos Results: Analysis of c-Fos expression

To test the prediction that the retrieval of generalized memory is increasingly supported by medial prefrontal cortical regions, while retrieval of the context-specific memory engages the hippocampus, we analyzed expression of c-Fos in the hippocampus and the aCC (Figure 1C, 1E). To assess c-Fos expression in the hippocampus, a 2 × 2 ANOVA was conducted with Context and Delay as between-subject factors. A significant main effect was found for Context, with higher c-Fos expression following context memory testing in CXT-A than in CXT-B (F(_1,23_) = 10.758, p = 0.004, η^2^_p_ = 0.350), but a non-significant main effect of Delay (F(_1,23_) = 0.327, p = 0.574, η^2^_p_ = 0.016) and a non-significant Context × Delay interaction (F(_1,23_) = 0.727 p = 0.404, η^2^_p_ = 0.035; Figure 1D). A similar pattern of results was found in both the dorsal and ventral hippocampus (data not shown, but available upon request).

To assess c-Fos expression in the aCC, a 2 × 2 ANOVA was conducted with Context and Delay as between-subject factors. A significant main effect was found for Delay, with higher c-Fos expression at memory retrieval in the LD condition (F(_1,28_) = 5.789, p = 0.024, n^2^_p_ = 0.188), but a non-significant main effect of Context (F(_1,28_) = 1.654, p = 0.210, n^2^_p_ = 0.062) and no significant Context x Delay interaction (F(_1,28_) = 0.146 p = 0.705, n^2^p = 0.006, Figure 1F).

Together, these results support the time-dependent shift in retrieval-associated activity within the hippocampus and mPFC nodes of the context fear memory network. The hippocampus is sensitive to context-specificity at both short and long delays, whereas it is primarily the prefrontal cortex that mediates the time-dependent generalization of the memory across contexts. Contrary to the traditional view of consolidation, these results suggest that the memory supported by the prefrontal cortex at long delays is qualitatively different (more generalized) from the memory that strongly engages the hippocampus. These results also suggest that although the hippocampus continues to be recruited for context-specific remote memory retrieval, a reorganized memory trace forms in the medial prefrontal cortex over time, and it is the latter that dominates behavioural performance at remote intervals.

**Figure 1.**
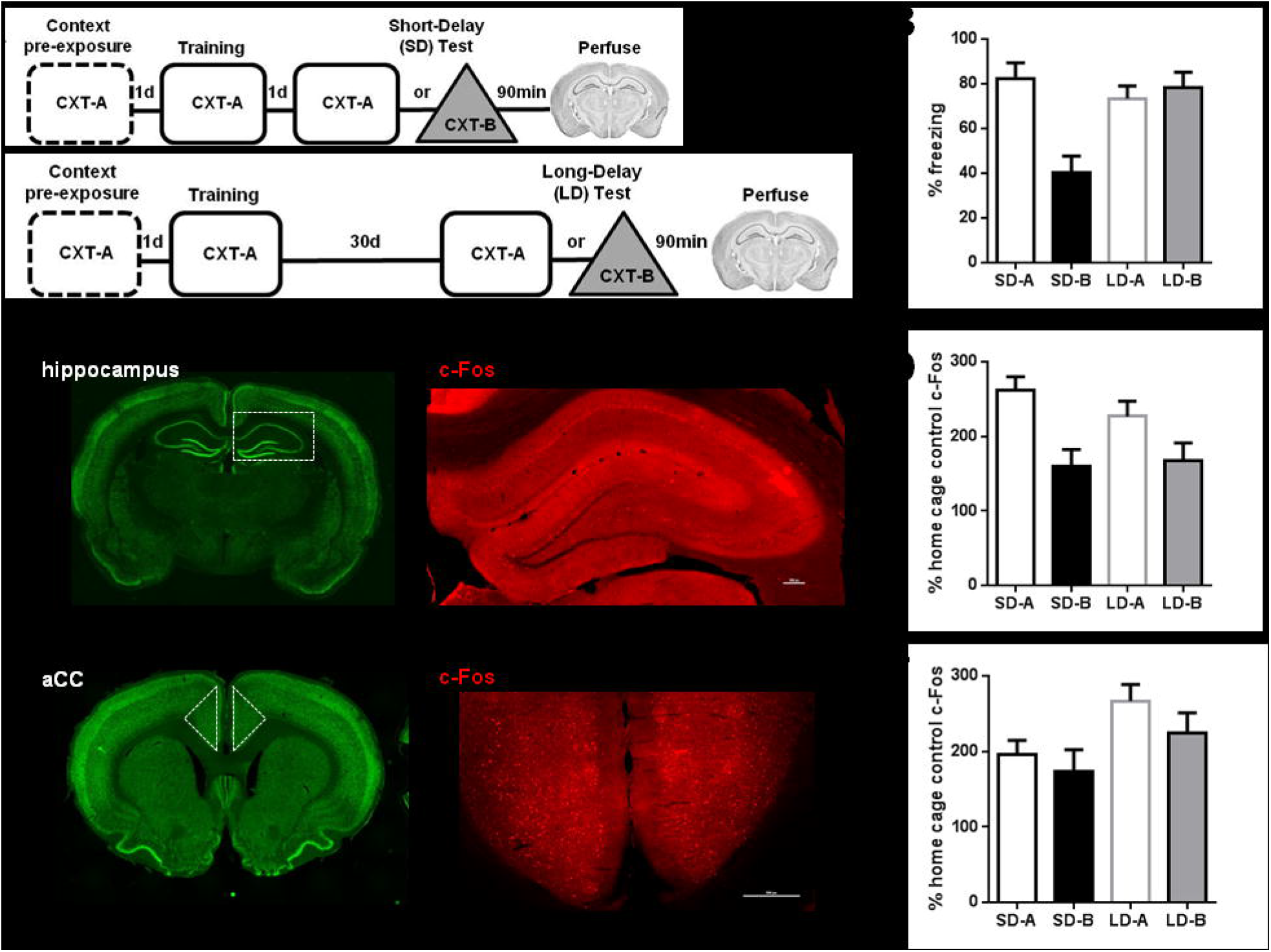
The time-dependent generalization of context memory and changes in cFos expression. A) Experimental timeline for the Short Delay (top) and for the Long Delay conditions (bottom) in Experiment 1. B) Mean percent time spent freezing during the first 6 min of the context fear memory test. Error bars represent the SEM. Rats froze significantly more in CXT-A than in CXT-B at the Short-Delay, but froze at similarly high levels in both CXT-A and CXT-B at the Long-Delay. C) Left: Coronal sections identifying the hippocampus (outlined in white) stained with NeuN (green) to label all neuronal nuclei. Right: Example of c-Fos staining (red) in the hippocampus. D) c-Fos expression levels in the hippocampus were significantly higher when tested in CXT-A than in CXT-B. Hippocampal c-Fos expression levels in each context did not differ between the Short and Long-Delay tests. Values are expressed as a percent change from the home cage control baseline c-Fos level. E) Left: Coronal sections identifying the aCC (outlined in white) stained with NeuN (green) to label all neuronal nuclei. Right: Example of c-Fos staining (red) in the aCC. F) c-Fos expression levels in the aCC were significantly higher when tested after a Long-Delay than a Short-Delay. aCC c-Fos expression levels did not differ between the CXT-A and CXT-B test conditions. Values are expressed as a percent change from the home cage control baseline c-Fos level. Error bars represent the SEM.

### Experiment 2: The neural basis of episodic memory transformation in humans

#### Behavioral Results In-scanner retrieval

Memory for short film clips was tested during fMRI scanning either immediately following encoding (0d), or after a delay of one week (7d). Significantly more film clips were forgotten during the 7d retrieval session compared to immediate (0d) retrieval (t(_19_) = −8.718, p <.0001) (Figure 2D), with a range of 0 to 5 forgotten clips during 0d retrieval, and a range of 1 to 11 forgotten clips during 7d retrieval.

A 2 × 2 repeated measures ANOVA was conducted with Rating Type (story content and vividness) and Delay (0d, 7d) as within-subject factors, and the assigned ratings as the dependent variable. The ANOVA revealed significant main effects of Rating Type (F(_1,19_) = 73.908, p < 0.0001, n ^2^_p_ = 0.795), with participants giving higher ratings for story content over vividness, and of Delay (F(_1,19_) = 55.555, p < 0.0001, n ^2^_p_ = 0.745), with a decrease in memory ratings over time. There was no significant interaction between Rating Type × Delay (F(_1,19_) = 0.209, p = 0.653, n^2^_p_ = 0.001; Figure 2C, left). This was confirmed by a paired-samples t-test for the percentage of ratings retained over time. The t-test revealed no significant difference between the percentage of story content or perceptual vividness ratings between the immediate and 7d retrieval sessions (t_(19)_ = 1.167, p = 0.257; Figure 2C, right).Together, these results suggest that participants judged that their memory declined equally over time for both main story content as well as the accompanying perceptual detail.

#### Post-scan retrieval

The qualitative content of the memories was evaluated immediately following each scanning session. Consistent with our previous work (Sekeres et al., 2016), participants demonstrated a greater loss of peripheral details (indicative of perceptual information) than of central elements (reflecting the schematic story content) over time. A 2 × 2 repeated measures ANOVA was conducted with Detail Type (central and peripheral) and Delay (0d, 7d) as within-subject factors, and the number of correctly recalled details (Retrieval Success) as the dependent variable. The ANOVA revealed significant main effects of Detail Type (F(_1,18_) = 57.977, p < 0.0001, η^2^_p_ = 0.763), with more peripheral details recalled than central elements, and of Delay (F(_1,18_) = 75.844, p < 0.0001, η^2^_p_ = 0.808), indicating that participants recalled fewer details after 7d. A significant Detail Type × Delay interaction (F(_1,18_) = 48.723, p < 0.0001, η^2^_p_ = 0.730) showed that memory for peripheral details suffered a significantly greater decline over time than memory for central elements (Figure 2E, left). Given that participants recalled central details near ceiling levels during 0d retrieval, we next confirmed that the differential rates of forgetting were not due to the different maximal number of retrievable peripheral details and central elements by conducting an additional analysis using percentage of details retained as the dependent variable. Consistent with the previous result, this analysis revealed a significantly greater percentage of retained central elements than of peripheral details over a week’s delay (t(_18_) = 4.735, p < 0.001; Figure 2E, right). These results indicate that memory for peripheral details declined disproportionally over the week following encoding, whereas memory for central story elements was preferentially retained. Of interest, this pattern was not reflected in subjects’ vividness ratings, as they reported equivalent retention of both story (central) and perceptual (peripheral) content.

**Figure 2.**
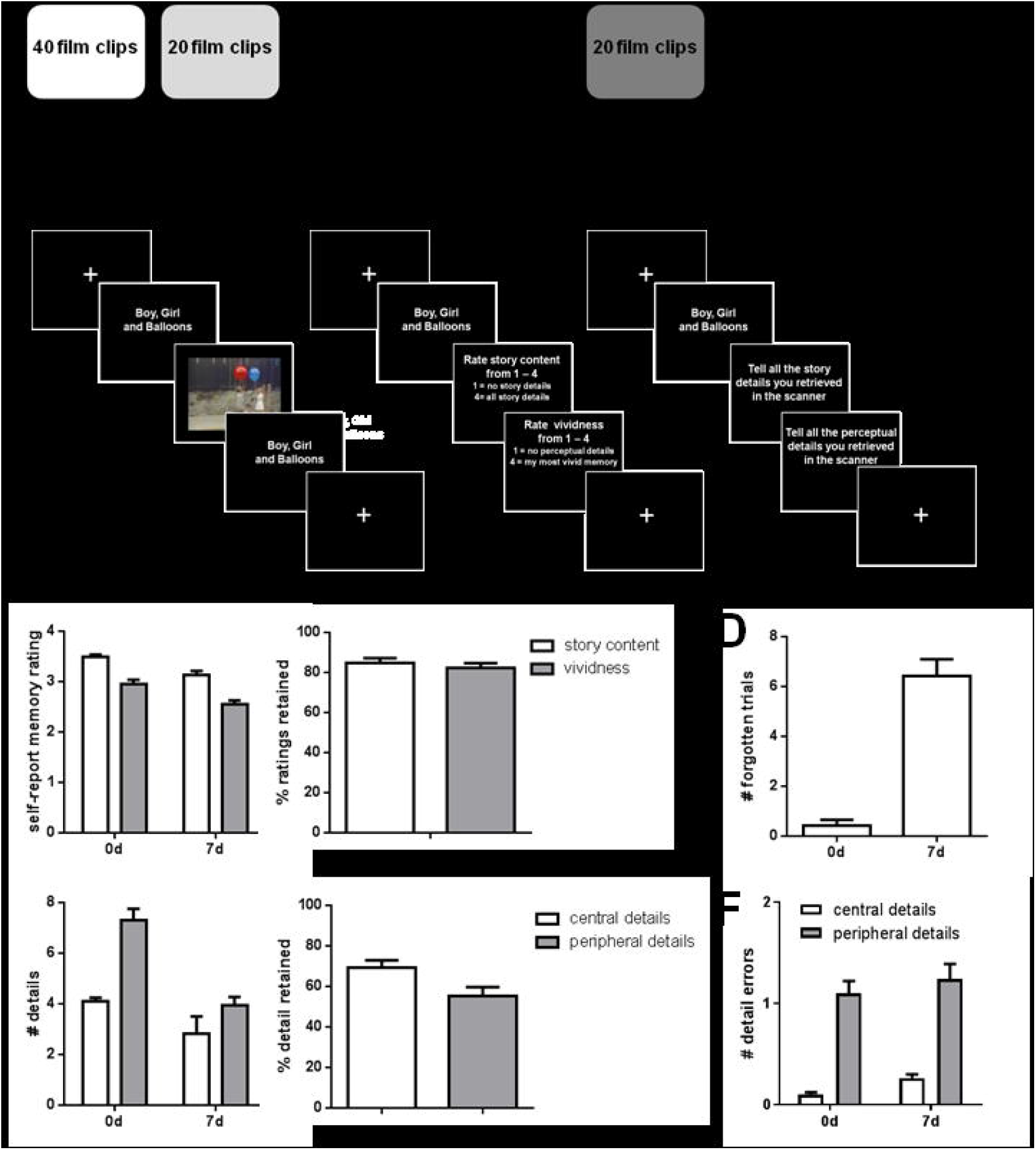
Time-dependent schematization of episodic memory. A) Experimental timeline for Experiment 2. B) Detailed schematic of the study design for the encoding session (left), in-scan retrieval session (middle), and post-scan retrieval session (right). Encoding session: 40 film clips were shown to participants in a randomized order. Retrieval session: The retrieval sessions were identically run across each delay (0d, 7d). C) Left: In-scanner memory ratings of the story content (white bars) and the vividness of perceptual details (grey bars) for each memory retrieval test session (0d and 7d delay). Participants rated their memory for the story content more highly than the vividness of their memory for the perceptual details in the film clips. Right: Percent of memory ratings retained between the 0d and 7d in-scanner retrieval sessions. The memory ratings declined at similar levels for both story content and vividness of perceptual details over the course of one week. D) Mean number of forgotten trials at the 0d and 7d delay. Forgotten trials were classified based on in-scanner memory ratings of 1 (lowest rating) for both story content and perceptual content. E) Left: Number of details (central elements and peripheral details) reported during the post-scanner verbal memory retrieval test session at each delay. Participants reported more peripheral details (grey bars) than central elements (white bars) when tested immediately after encoding but similar levels of central elements and peripheral details when tested 7d following encoding. Right: Percent of memory details retained between the 0d and 7d post-scanner retrieval sessions. Over the course of one week, peripheral details were forgotten at a higher rate than central elements. F) Mean number of errors per trials during the 0d and 7d post-scan verbal retrieval. Errors were subtracted from the total number of retrieved details to produce the corrected number of central and peripheral details at each delay (Retrieval Success). Error bars represent the SEM.

To assess changes in errors during memory retrieval, A 2 × 2 repeated measures ANOVA was conducted with Detail Type (central and peripheral) and Delay (0d, 7d) as within-subject factors. The ANOVA revealed significant main effects of Detail Type (F(_118_) = 81.133, p < 0.0001, n^2^_p_ = 0.818) with more errors in peripheral details than central elements, and of Delay (F(_1_,_18_) = 4.724, p = 0.043, n^2^_p_ = 0.208), indicating that participants made more errors during the 7d retrieval session. There was no significant Detail Type x Delay interaction (F(_1,18_) = 0.036, p = 0.852, n^2^_p_ = 0.002; Figure 2F). Taken together, although there was a time-dependent increase in errors, and a higher incidence of errors in peripheral details, these errors were accounted for when calculating the corrected ‘Retrieval Success’ score.

Analyses were also conducted to determine any gender differences in ratings and detailed memory retrieval using ANOVA with gender as a between-subject factor. We found a trending main effect of gender for ratings (F(_1,18_) = 4.072, p =0.059, n^2^_p_ = 0.184) with males rating the quality of their memory retrieval higher than did females, but no significant main effects or interactions for any other analyses (all p >0.1).

#### fMRI Results: Analysis of BOLD activity

To test the prediction that recent, perceptually detailed memories for the film clips are supported by hippocampal activity, whereas older memories, being less perceptually detailed, yet retaining most central elements, are supported by medial prefrontal cortical regions, we assessed modulations of fMRI BOLD activity across the brain during film clip retrieval using Partial Least Squares (PLS). This multivariate approach to assessing co-varying patterns of activity is more sensitive than univariate approaches (Fletcher et al., 1996; Lukic et al., 2002), and allows us to take full advantage of fMRI’s ability to identify brain-wide retrieval networks including and extending beyond the hippocampal and medial prefrontal cortical regions at each retrieval delay. To identify patterns of retrieval activity that characterized each delay, we first contrasted memory retrieval at each delay separately with a fixation control task. The significant increases in activity (p < 0.001) during immediate (0d) retrieval are shown in warm colors in Figure 3A, and positive brain scores in Figure 3B. Immediate retrieval activated the middle and inferior temporal gyri, as well the medial temporal lobe, including the right parahippocampal cortex (cluster includes activity in the right anterior and posterior hippocampus), and the left fusiform gyrus (cluster includes activity in the left anterior and posterior hippocampus, and parahippocampal cortex). During immediate retrieval, activity also was evident in the prefrontal cortex, including clusters in the left inferior frontal gyrus and insular cortex, the right precentral gyrus, as well as the bilateral precuneus (See Table 1 (top) for full list of regions).

We next contrasted 7d delayed retrieval with fixation. The significant activations above control activity (p < 0.001) seen at 7d retrieval are shown in warm colors in Figure 3C, and positive brain scores in Figure 3D. Memory retrieval 7d after encoding was accompanied by activation in the left superior temporal gyrus and bilateral middle and inferior temporal gyri, as well as the medial temporal lobe, including the right anterior hippocampus and fusiform gyrus (cluster includes the right parahippocampal cortex), and the left fusiform gyrus (cluster includes the left hippocampus, and parahippocampal cortex). Distributed activity was also seen in the prefrontal cortex, including large clusters in the right superior frontal gyrus, bilateral middle and inferior frontal gyrus, bilateral insular cortex, as well as the right precuneus (See Table 1, bottom, for full list of regions).

To determine differences in the patterns of activity elicited during immediate and 7d delayed memory retrieval, we directly contrasted retrieval-related activation at each delay. A significant pattern (p = 0.008) differentiating retrieval at the two delays showed more activity for immediate memory in the right posterior hippocampus and the bilateral precentral gyrus (Figure 3E warm colored regions, and positive brain scores in Figure 3F). Relative to 0d, 7d memory retrieval (Figure 3E cool colored regions, and negative brain scores in Figure 3F) activated the bilateral left middle frontal gyrus, inferior frontal gyrus and medial prefrontal cortex, including the bilateral superior medial frontal gyrus, as well as the right aCC (See Table 2 for full list of regions).

Together, these three PLS analyses identify retrieval-related activity in medial temporal and prefrontal cortical regions that is present across delays (when contrasted with fixation), and highlight a shift in the recruitment of these regions. Specifically, hippocampal and parahippocampal activity declines, but does not completely disappear, as activity in the medial and lateral prefrontal cortex increases over time. This shift in relative hippocampal and prefrontal activity as the memory ages and loses a disproportionate amount of peripheral detail mirrors the time-dependent pattern seen in rodents, where activity shifts towards the aCC as remote memory generalizes across contexts (Experiment 1).

**Figure 3.**
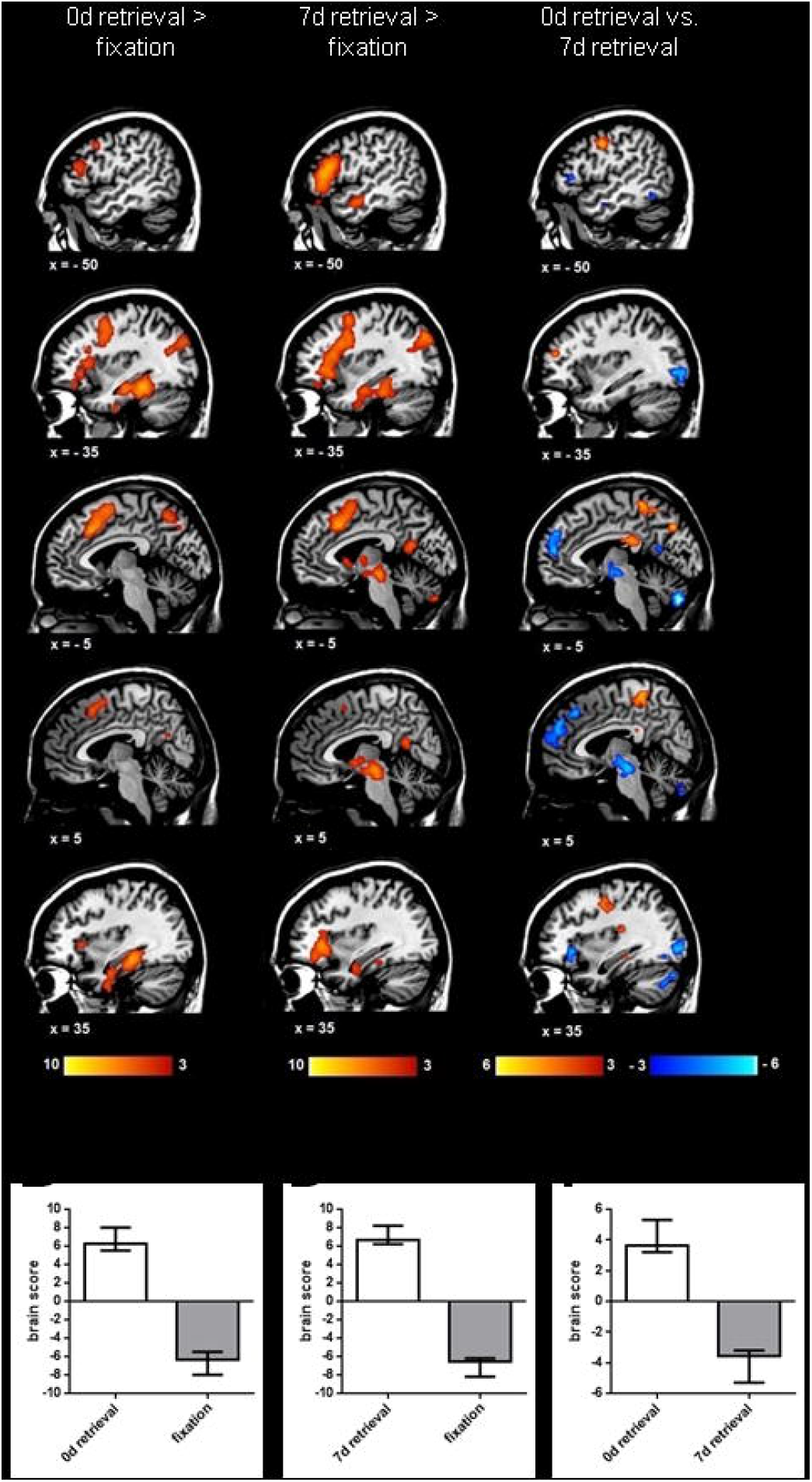
Time-dependent changes in the network of brain regions active during memory retrieval. Mean-centered, blocked design PLS analyses were conducted to assess modulations in BOLD activity across the brain during memory retrieval. A) LV depicting brain activity associated with immediate (0d) retrieval (warm colors) contrasted with fixation (areas with more activity during fixation are not shown on figure). Note the anterior and posterior hippocampal activation during the immediate retrieval task. B) Brain scores reflecting the degree to which 0d retrieval (positive BSRs) and fixation (negative BSRs) correlate with the pattern of activity seen in 3A. Error bars are 95% confidence intervals. C) LV depicting brain activity associated with 7d delayed retrieval (warm colors) contrasted with fixation (data not shown on figure). Note the less extensive posterior hippocampal activation, and the increased medial prefrontal activation during the 7d retrieval task. D) Brain scores reflecting the degree to which 7d retrieval (positive BSRs) and fixation (negative BSRs) correlate with the pattern of activity in 3C. Error bars are 95% confidence intervals. E) LV depicting brain activity associated with immediate (0d) retrieval (warm colors) contrasted with 7d delayed retrieval (cool colors). F) Brain scores reflecting the degree to which 0d retrieval (positive BSRs) and the 7d retrieval (negative BSRs) correlate with the pattern of activity in 3E. Error bars are 95% confidence intervals. fMRI results are displayed using Mango (Research Imaging Institute, UTHSCSA). PLS = Partial Least Squares; BOLD = blood-oxygen-level dependent; LV = latent variable; BSR = bootstrap ratio.

**Table 1.**
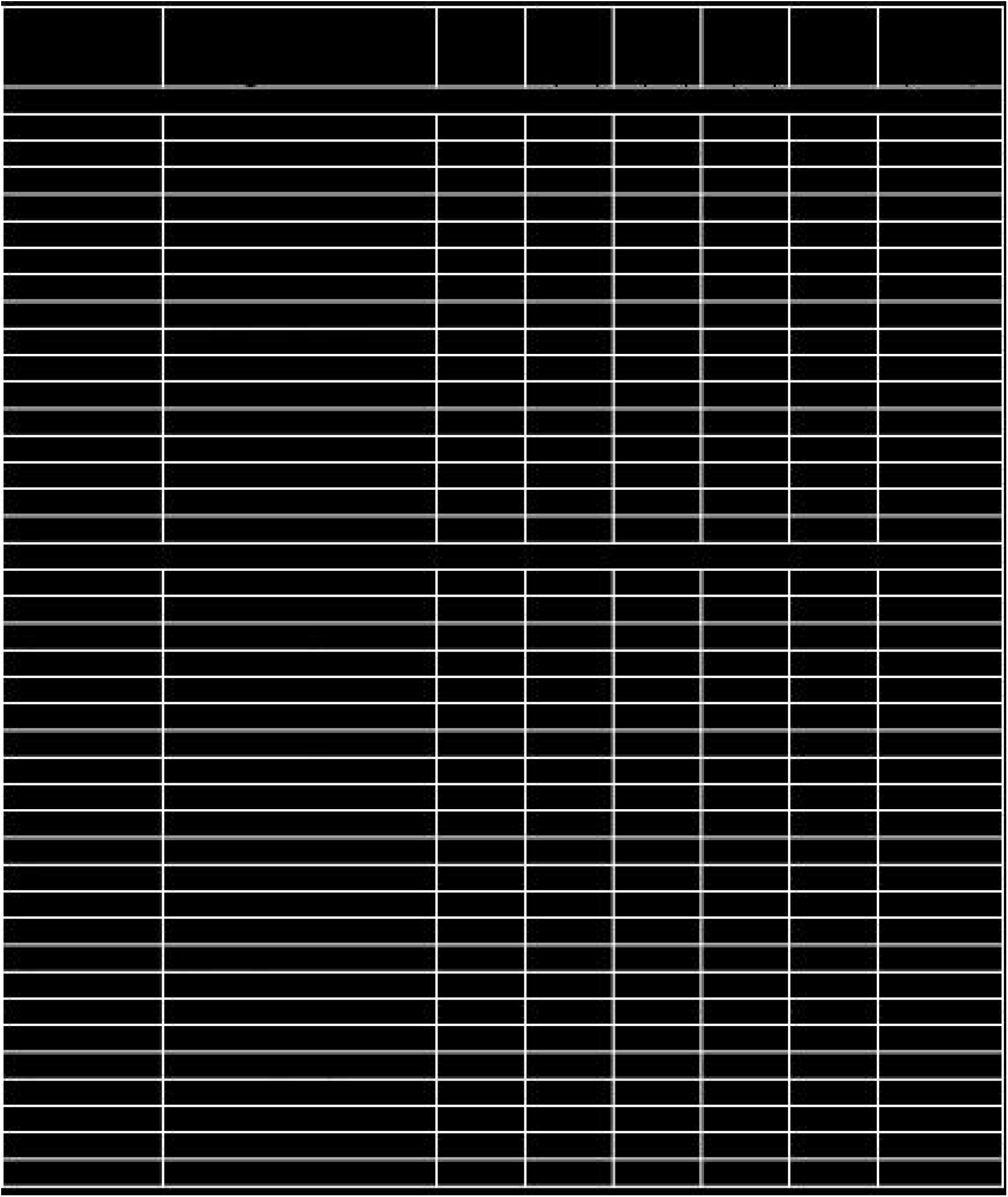
Coordinates of regions associated with memory retrieval. Top: MNI coordinates of the peak activation voxel within each cluster are reported for the contrast of immediate (0d) retrieval to fixation (warm color activations in Figure 3A, positive BSRs, data for fixation > retrieval are not shown in Figure 3A). Bottom: MNI coordinates of the peak activation voxel within each cluster are reported for the comparison of 7d retrieval to fixation (warm color activations in Figure 3C, positive BSRs, data for fixation > retrieval are not shown in Figure 3C). Hem = hemisphere; BA = Brodmann Area; BSR = bootstrap ratio from the PLS analysis indicating the robust contribution of the reported voxel. *indicates the cluster contains structures within the retrieval network.

**Table 2.**
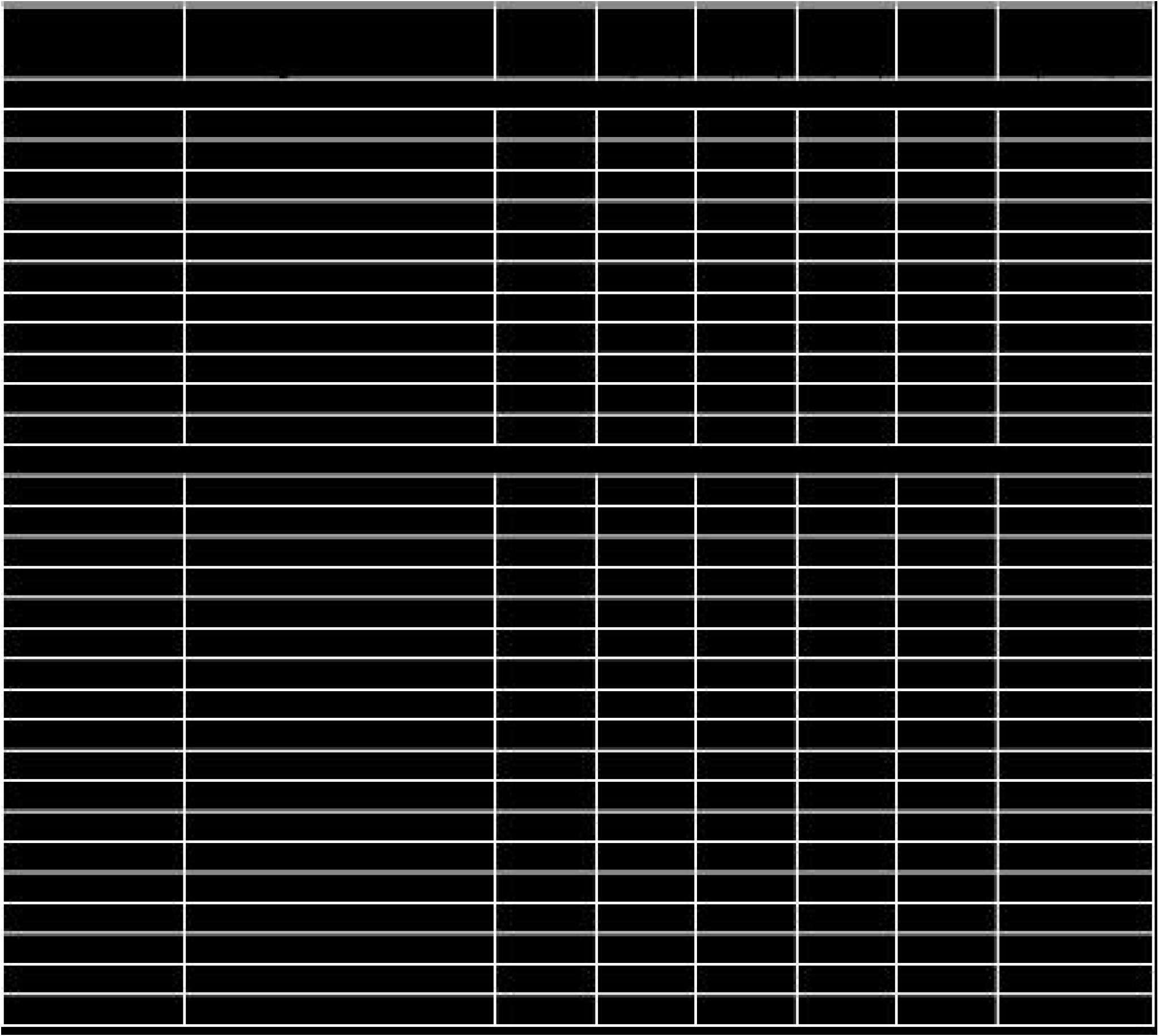
Coordinates of regions associated with immediate and 7d delayed retrieval. MNI coordinates of the peak activation voxel within each cluster are reported for LV1 for the PLS analysis of 0d retrieval (warm color activations in Figure 3E, positive BSRs) contrasted with 7d retrieval (cool color activations in Figure 3E, negative BSRs). Hem = hemisphere; BA = Brodmann Area; BSR = bootstrap ratio from the PLS analysis indicating the reliability of the reported voxel. *indicates the cluster contains structures within the retrieval network.

To determine if retrieval of detailed episodic memory continues to elicit hippocampal activity in humans, we contrasted the immediate retrieval session and the 7d delayed retrieval session only for clips that were rated as vividly retrieved (clips given in-scan perceptual vividness and story content ratings of 3 or 4). The in-scanner ratings, rather than the detail scores, were used to classify clips due to the fact that the ratings were a measure of performance taken immediately after each in-scanner retrieval trial. In this analysis, if older but vividly recalled memories continue to engage the hippocampus, we would not expect to see differences in hippocampal activity between these two conditions. As predicted, no delay-related differences in hippocampal or parahippocampal activity were found for vivid retrieval, though differences were evident in other brain regions. We identified a significant distributed pattern of activity (p =0.017) that differentiated vivid memory retrieval between the two delays. This pattern indicated greater activity in a small cluster in the right insular cortex for immediate than for 7d old vivid memory (Figure 4A warm colored regions, and positive brain scores in Figure 4B). Relative to 0d, 7d vivid memory retrieval (Figure 4A cool colored regions, and negative brain scores in Figure 4B) activated bilateral clusters in the inferior frontal gyrus and medial prefrontal cortex including the left superior medial frontal gyrus and the right aCC. Activity was also observed in the pCC and angular gyrus (see Table 3 for full list of regions).

**Figure 4.**
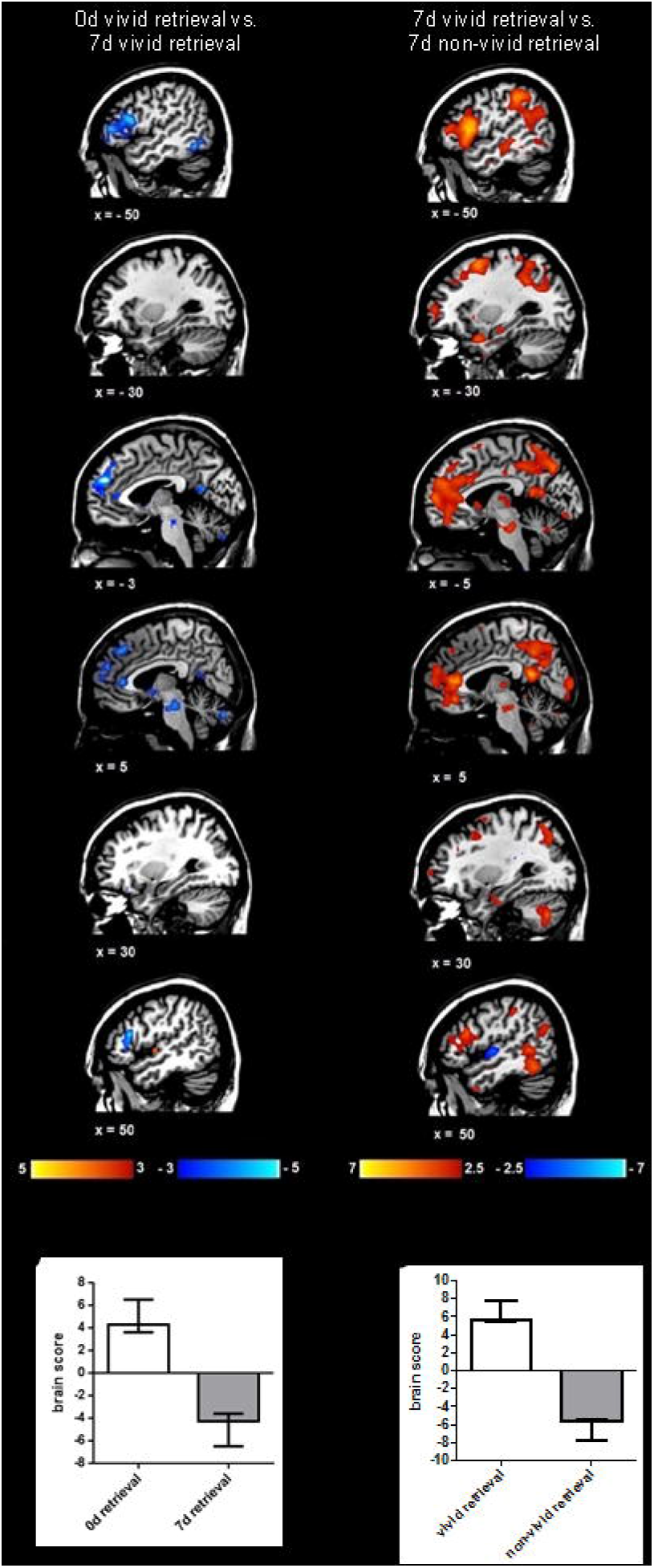
Vividly retrieved film clips do not differ in hippocampal activity during immediate and 7d delayed retrieval. A) LV depicting brain activity associated with immediate (0d) vivid retrieval (warm colors) contrasted with 7d delayed vivid retrieval (cool colors). No differences were observed in hippocampal activation between the 0d and 7d retrieval session. There is increased medial prefrontal activation, and pCC and angular gyrus activity accompanying memory vivid retrieval after 7d. B) Brain scores reflecting the degree to which 0d vivid retrieval (positive BSRs) and 7d vivid retrieval (negative BSRS) correlate with the pattern of activity seen in 4A. C) LV depicting brain activity associated with 7d vivid retrieval (warm colors) contrasted with 7d non-vivid retrieval (cool colors). Note the bilateral hippocampal activity observed during vivid 7d retrieval. D) Brain scores reflecting the degree to which 7d vivid retrieval (positive BSRs) and 7d non-vivid retrieval (negative BSRs) correlate with the pattern of activity in 4E. Error bars are 95% confidence intervals.

**Table 3.**
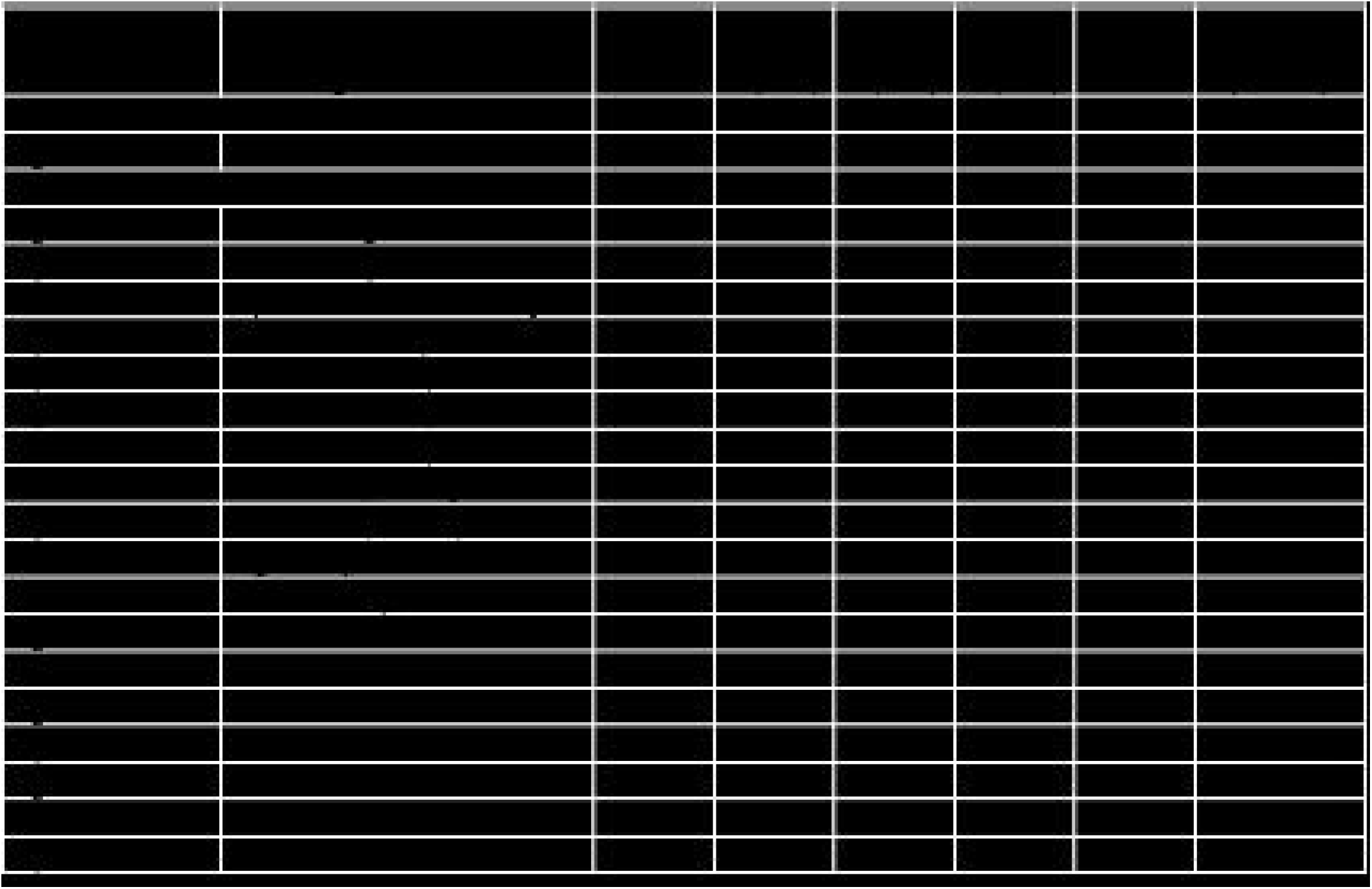
Coordinates of regions associated with immediate and 7d delayed vivid retrieval. MNI coordinates of the peak activation voxel within each cluster are reported for LV1 for the PLS analysis of 0d vivid retrieval (warm color activations in Figure 4A, positive BSRs) contrasted with 7d vivid retrieval (cool color activations in Figure 4A, negative BSRs). Hem = hemisphere; BA = Brodmann Area; BSR = bootstrap ratio from the PLS analysis indicating the reliability of the reported voxel. *indicates the cluster contains structures within the retrieval network.

The absence of a significant difference in hippocampal activity between 0 and 7d retrieval for vivid memories is consistent with the hypothesis that for vivid memories, the hippocampus is not differentially engaged at different delays. It should be noted, however, that the 7d vivid retrieval analysis contains fewer trials than used in the previously reported analyses, and the failure to detect differences in hippocampal activation may, in part, be a result of low power. Therefore, to confirm the continuing role of the hippocampus during vivid memory retrieval after 7d, we contrasted vivid retrieval with non-vivid retrieval during the 7d session. The significant increases in activity (p < 0.001) seen during 7d vivid retrieval are shown in warm colors in Figure 4C and positive brain scores in Figure 4D. Significant increases in activity during 7d non-vivid retrieval are shown in cool colors, and negative brain scores. Vivid retrieval activated clusters in the medial temporal lobe, including bilateral hippocampus, and parahippocampal cortex. These results confirmed robust hippocampal activity during retrieval of highly vivid memories, but not for those equally aged memories retrieved with low vividness.

In line with the prediction that memories engage a changing network of brain regions over time, vivid retrieval of clips after 7d, compared to non-vivid retrieval, was accompanied by additional activation of clusters in the medial and lateral frontal lobes, as well as in other regions typically involved in the retrieval network, including the middle temporal gyrus, bilateral pCC, precuneus, angular gyrus, and areas of the ventromedial prefrontal cortex. These results suggest a time-dependent broadening of the network supporting vivid memory retrieval that involves a shift towards increased recruitment of frontal nodes in the network (See Table 4 for full list of regions). Increased activity associated with 7d non-vivid retrieval was found in small clusters in the right superior temporal gyrus and caudate.

**Table 4.**
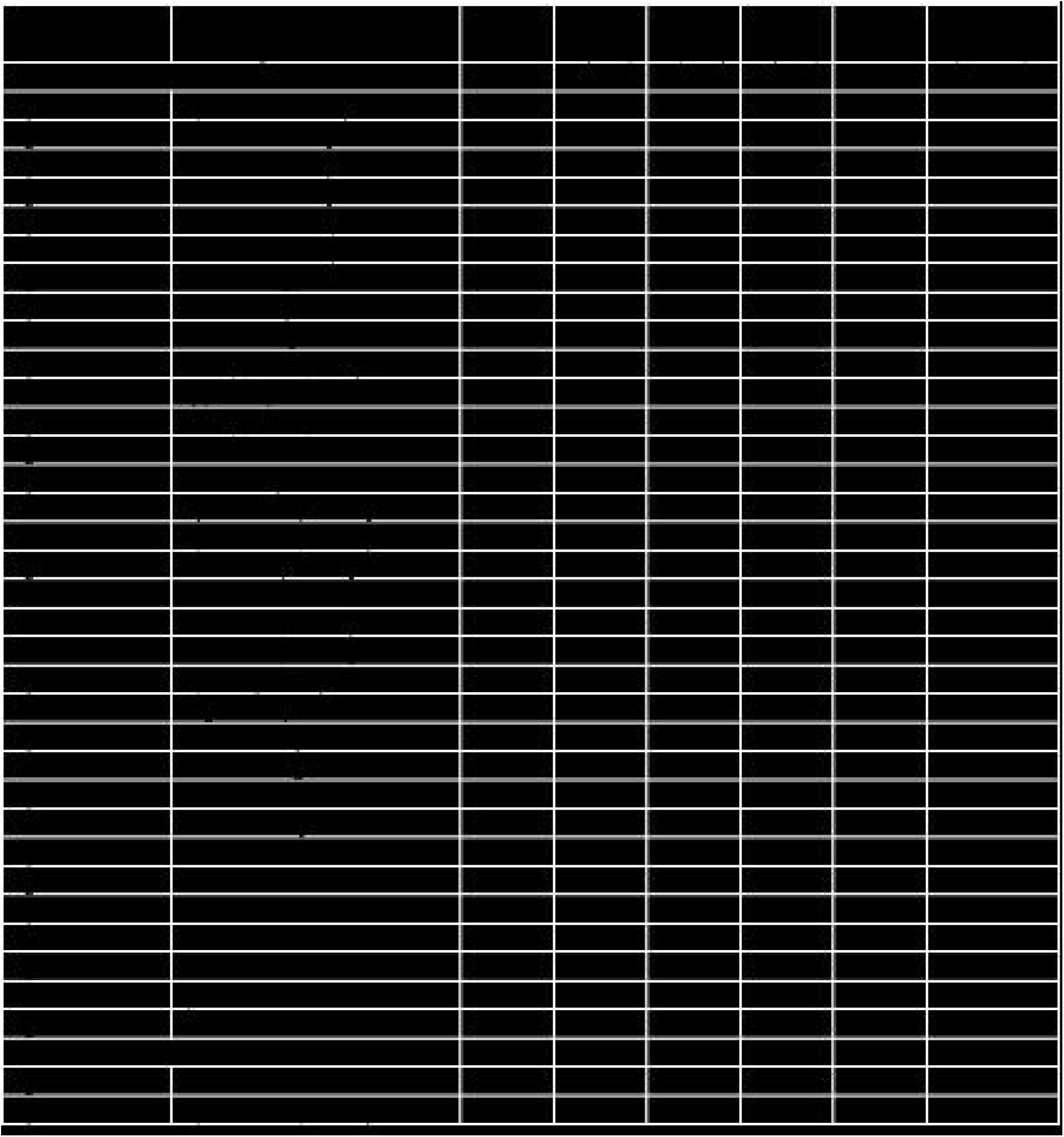
Coordinates of regions associated with 7d vivid and non-vivid memory retrieval. MNI coordinates of the peak activation voxel within each cluster are reported for LV1 for the PLS analysis of 7d vivid retrieval (warm color activations in Figure 4C, positive BSRs) contrasted with 7d non-vivid retrieval (cool color activations in Figure 4C, negative BSRs). Hem = hemisphere; BA = Brodmann Area; BSR = bootstrap ratio from the PLS analysis indicating the robust contribution of the reported voxel. *indicates the clusters falls within the retrieval network.

## Discussion

Using complementary neuroimaging approaches in rodents and humans we show time-dependent changes in the quality of retrieved memories corresponding to changes in hippocampal-mPFC activity. Our results show that (1) the hippocampus is strongly activated during retrieval of context-specific memory in rats, and of detailed episodic memory in humans; (2) retrieval of general or schematic memory is supported by increased mPFC activity, and reduced hippocampal activity; (3) older memories that retain context-specificity (in rodents) or perceptual detail (in humans) continue to engage the hippocampus, yet increasingly recruit mPFC. These observations across species support the idea of a dynamic interplay between hippocampal and prefrontal cortical regions as memories transform over time, and suggest this interplay is influenced by both the age and the nature of the retrieved memory.

### The hippocampus continues to support context-specific and episodic memory

In rodents, the hippocampus was more active when memory is tested in the original conditioning context than in a novel context, regardless of the memory’s age (Wiltgen et al., 2010). Optogenetic studies report that hippocampal neuronal ensembles engaged during memory acquisition continue to support context-specific memories. Although, over time, the network may distribute in the cortex, activation of the original cell assembly can induce expression of the context memory (Liu et al., 2012; Ramirez et al., 2013), while rapid inactivation leads to memory loss (Goshen et al., 2011). Likewise, in humans, 7d vivid memory retrieval robustly engaged the hippocampus. The lack of difference in hippocampal activation between vivid retrieval at 0d and 7d was likely not due to the limited clips that were vividly retrieved after 7d, because retrieval of 7d-old vivid memories shows strong hippocampal activity when contrasted with non-vivid retrieval at that same timepoint.

Together, these findings support the notion that the *nature*, rather than the age, of a retrieved memory determines hippocampal recruitment in both species. If the memory remains detailed, vivid (humans) and context-specific (rodents), the hippocampus continues to support its representation (Winocur and Moscovitch, 2011) and does not disengage over time (Squire and Bailey, 2007).

### Medial prefrontal cortex activity Increases as memories lose specificity

In rodents, mPFC becomes increasingly active during remote context memory retrieval (Frankland et al., 2004; Wheeler et al., 2013). Critically, we demonstrate that, unlike hippocampus, mPFC activity is relatively insensitive to the retrieval context. This finding helps resolve a long-standing debate concerning the nature of this representation. We suggest that a memory which becomes represented in mPFC is not an identical copy of the original memory, but rather a transformed, less detailed and vivid version that is qualitatively different from one which engages both regions (Winocur et al., 2010). Evidence from hippocampal lesion or inactivation studies supports this interpretation (Winocur et al., 2007; Denny et al., 2014; Einarsson et al., 2014; Cullen et al., 2015).

In Experiment 2, the relative contributions of hippocampus and mPFC were mediated by the memory’s age and qualitative content. Immediately following encoding, memories contained most of the central details defining the film’s events, and peripheral (perceptual, contextual) details. Retrieval was supported by strong, bilateral activity in anterior and posterior hippocampus. After 7d, many peripheral details were forgotten, while central elements were relatively preserved. Retrieval of this less-detailed, schematic memory was supported by increased activity in mPFC, and continued activation of anterior hippocampus. These findings are consistent with reports of functional specialization along the long-axis of the hippocampus, and its mPFC projections. Anterior hippocampus codes global representations related to the central elements of an event (Poppenk and Moscovitch, 2011; Poppenk et al., 2013; Robin and Moscovitch, 2017). Accordingly, persistent activity in the anterior hippocampus during 7d retrieval can account for both the increase in mPFC activity, and retrieval of the more schematic version of the memory (Ghosh et al., 2014). Reduced posterior hippocampal activity, and the decline of peripheral details and vividness after 7d is consistent with the proposal that posterior hippocampus codes for finely detailed representations of an event (Moscovitch et al., 2016).

Together, the results of both experiments are consistent with TTT, which proposes that a memory undergoes a transformation during which its schematic features are represented cortically, whereas both fine and coarse contextual and perceptual details characterizing the original experience continue to be represented in the hippocampus (Nadel and Moscovitch, 1997; Moscovitch et al., 2016). To retrieve these details, the hippocampus remains necessary, regardless of the memory’s age. Contextual cues prior to retrieval can reactivate the context-specific memory, and reinstate hippocampal dependency in rodents (Winocur et al., 2009) and humans (Cohn et al., 2009), suggesting that both the generalized and detailed versions of memories can co-exist, with the conditions at the time of retrieval determining which version will be expressed.

A caveat to consider when using context fear conditioning to understand memory consolidation is the inclusion of aversive associative memories, which likely engages additional fear memory neural systems. However, earlier investigations report a similar shift towards mPFC activity as spatial memories age (Bontempi et al., 2000; Richards et al., 2014), suggesting a common transformation process operating upon hippocampal-dependent memories (Winocur et al., 2005, 2009).

Loss-of-function studies have confirmed the necessity of the hippocampus in context-specific memories in rodents and perceptually-detailed episodic memories in humans (Winocur and Moscovitch, 2011). Correlational studies such as the present ones are important for understanding the recruitment of regions within the retrieval network under normal physiological conditions in order to better understand memory network dynamics. The present experiments tested for the continuing recruitment of the hippocampus over relatively short delays following memory encoding (7d in humans, 30d in rodents). While reduced hippocampal activity was observed as memories aged and lost detail, or generalized to other contexts, they did not completely disengage from the hippocampus, suggesting that, if intact, the hippocampus continues to participate in memory retrieval.

Given the current design, we cannot definitively state that the pattern of activity would be similarly observed after a period of months or years, although there is strong evidence for continuing hippocampal engagement during retrieval of decade old episodic memories (Bonnici and Maguire, 2017). While the timeline for memory transformation is prolonged, changes in the BOLD response during retrieval of different aged memories may reflect the early stages of this process. One week may seem a short period of time to expect large-scale changes in networks supporting memory retrieval in humans, but reorganization of declarative memory networks has been detected just 24-hrs following memory acquisition (Takashima et al., 2009; Ritchey et al., 2015). These findings suggest that reorganization of memory networks begins early, and may continue over the lifetime of a memory (Dudai et al., 2015).

### Reorganization of the memory network: Beyond hippocampus and mPFC

Investigation of whole-brain IEG activity following retrieval of recently acquired context memory in mice identified a network including hippocampus, medial temporal, and posterior parietal cortical regions (Vousden et al., 2015). These regions are similar to those we found active during recent memory retrieval in humans, suggesting that, despite noteworthy differences between animal and human memories, considerable overlap exists in the retrieval networks of both species. Chemogenetically silencing key hubs of the remote memory network in mice, including hippocampal CA1, reduced global efficiency of the network, and disrupted contextual fear memory (Vetere et al., 2017). This finding corresponds to similar deficits observed in humans after temporary inactivation of CA1 during transient global amnesia (Bartsch et al., 2011). How activity within the broader network in the rodent brain changes as a context-specific memory transforms over time remains unknown, but the results of Experiment 2 offer novel insight into this question.

In Experiment 2, immediate memory retrieval was supported by mPFC, hippocampus, parahippocampus, and posterior parietal activity. Not all regions of the retrieval network show significant activity above the fixation control task, likely due to overlap with the default mode network (Spreng et al., 2009; Spreng and Grady, 2010). Reduced activity throughout the posterior parts of this network (pCC, precuneus), was observed during 7d retrieval. Given the involvement of these regions in recollection, this result suggests that recollective processes are not likely engaged during retrieval of detail-poor memories (Svoboda et al., 2006; McDermott et al., 2009). Other key regions of the retrieval network showed robust activity when memories were vividly (relative to non-vividly) retrieved after 7d, including pCC and angular gyrus, areas typically associated with re-experiencing contextual and perceptual details during retrieval (Yonelinas, 2002). In turn, the shift towards prefrontal cortical activity within the network may reflect reliance on schematic knowledge (Gilboa and Marlatte, 2017), and the need for more attentional control and error monitoring (Moscovitch and Winocur, 2002; Gilboa et al., 2006; Cavanagh et al., 2009) during effortful retrieval of older memories.

Further investigations will determine the cellular mechanisms underlying memory transformation, how qualitative changes in memory are accompanied by shifts in functional connectivity between the hippocampus and mPFC, and changes in the weighting of key nodes in the retrieval network over time. Each of these avenues of research will require complementary contribution from studies of both humans and animals, with the ultimate goal of providing a neurobiological model of memory transformation across species.

## AUTHOR CONTRIBUTIONS

Conceptualization: MJS, GW, MM, CG; Methodology: MJS, GW, MM, JMW, MSL, MPM, CG; Investigation: MJS, JAEA, SP; Writing – Original Draft: MJS, GW, MM, CG; Writing – Review Editing: MJS, GW, MM, JAEA, SP, JMW, MSL, MPM, CG; Funding Acquisition: GW, MM, CG; Resources: JMW, MSL, GW, CG; Supervision: GW, MM, CG.

## Acknowledgements

We gratefully acknowledge the technical assistance of Jeremy Audia, Christy Cole-Harding, Christa Dang, Nick Hoang, Daniel Nichol, Yao-Fang Tan, Annette Weekes-Holder, and Marilyne Ziegler. Additional financial support came from the Canada Research Chairs program, the Ontario Research Fund, the Canadian Foundation for Innovation, and the Heart and Stroke Foundation Centre for Stroke Recovery. We would also like to acknowledge the generosity of Jack Anne Weinbaum, Sam Ida Ross, and Joseph Sandra Rotman in their support of the imaging centre at Baycrest.

## Notes

**Conflict of Interest**: The authors declare no competing financial interests

